# Influence of Lipomannan and Lipoarabinomannan Concentration on Mycobacterial Inner Membranes Characterized by All-atom Simulations

**DOI:** 10.64898/2026.04.15.718817

**Authors:** Hwayoung Lee, Nathaniel Rygh, Matthieu Chavent, Wonpil Im

## Abstract

Mycobacteria are responsible for causing severe illnesses like tuberculosis and leprosy in humans. Studying the mycobacteria cell envelope presents a significant challenge due to its intricate lipid compositions and structural variations and also its harmful nature in a typical experiment setting. In this study, we use all-atom molecular dynamics simulation to study mycobacterial inner membranes (MIMs). By incorporating different types of phosphatidyl-*myo*-inositol-mannosides (PIMs) and their glycoconjugates such as lipomannans (LM) and lipoarabinomannans (LAM) lipoglycans, we have constructed both symmetric and asymmetric membrane systems to study the MIM structure and dynamics under varying compositions of each lipid type. Our results show that the phospholipid/PIM-rich inner leaflet remains a stable, fluid bilayer, and the outer leaflet structure and dynamics are heavily governed by lipoglycan surface density. Importantly, as LM/LAM concentration increases, the polysaccharide chains shift from flexible, membrane-lying orientations to a compact brush-like state aligned with the membrane normal. This crowding significantly reduces the solvent-accessible volume and limits direct interactions between LM/LAM sugars and the outer leaflet surface. Furthermore, we observe that high lipoglycan presence in the outer leaflet slows lipid diffusion across the entire bilayer, demonstrating a dynamic coupling between the two leaflets. By resolving these LM/LAM sugar-level dynamics and their impact on membrane-wide properties, this study provides a molecular framework for future MIM modeling and simulation with various (peripheral) membrane proteins to better understand how the MIM functions as a regulated physical barrier and a platform for mycobacterial virulence.

## Introduction

The complex mycobacterial cell envelope is the defining feature of the genus and a major determinant of its physiology, pathogenicity, and intrinsic resistance to antibiotics. Rather than conforming to classical Gram-positive or Gram-negative paradigms, mycobacteria possess a highly specialized, multilayered envelope composed of an inner membrane, a thick and chemically diverse peptidoglycan-arabinogalactan layer, and an outer membrane enriched in mycolic acids^1–4^. This unusual architecture creates a formidable permeability barrier^5–7^ while simultaneously supporting a wide range of essential cellular processes. Despite its central importance, many aspects of the molecular organization and physical behavior of the mycobacterial inner membrane (MIM) remain poorly understood and have only recently begun to be explored using modern biophysical and computational approaches^8–10^.

A prominent characteristic of the MIM is the presence of complex glycoconjugates, among which lipomannan (LM) and lipoarabinomannan (LAM) lipoglycans are particularly abundant and biologically significant (**Figure 1**)^11–14^. Both molecules are anchored to the MIM through a phosphatidyl-*myo*-inositol-mannoside (PIM) moiety, while their extended polysaccharide domains project away from the membrane surface and into the periplasm. LM and LAM are involved in numerous aspects of mycobacterial biology, including modulation of membrane properties^15,16^, interactions with host immune cells^13,17^, and regulation of host–pathogen signaling pathways. LAM is well established as a key virulence factor capable of altering macrophage activation and phagosomal maturation^18–21^. Beyond their immunological roles, LM and LAM are the integral components of the MIM itself, yet their contributions to the MIM structure and dynamics are not well characterized.

**Figure 1.**
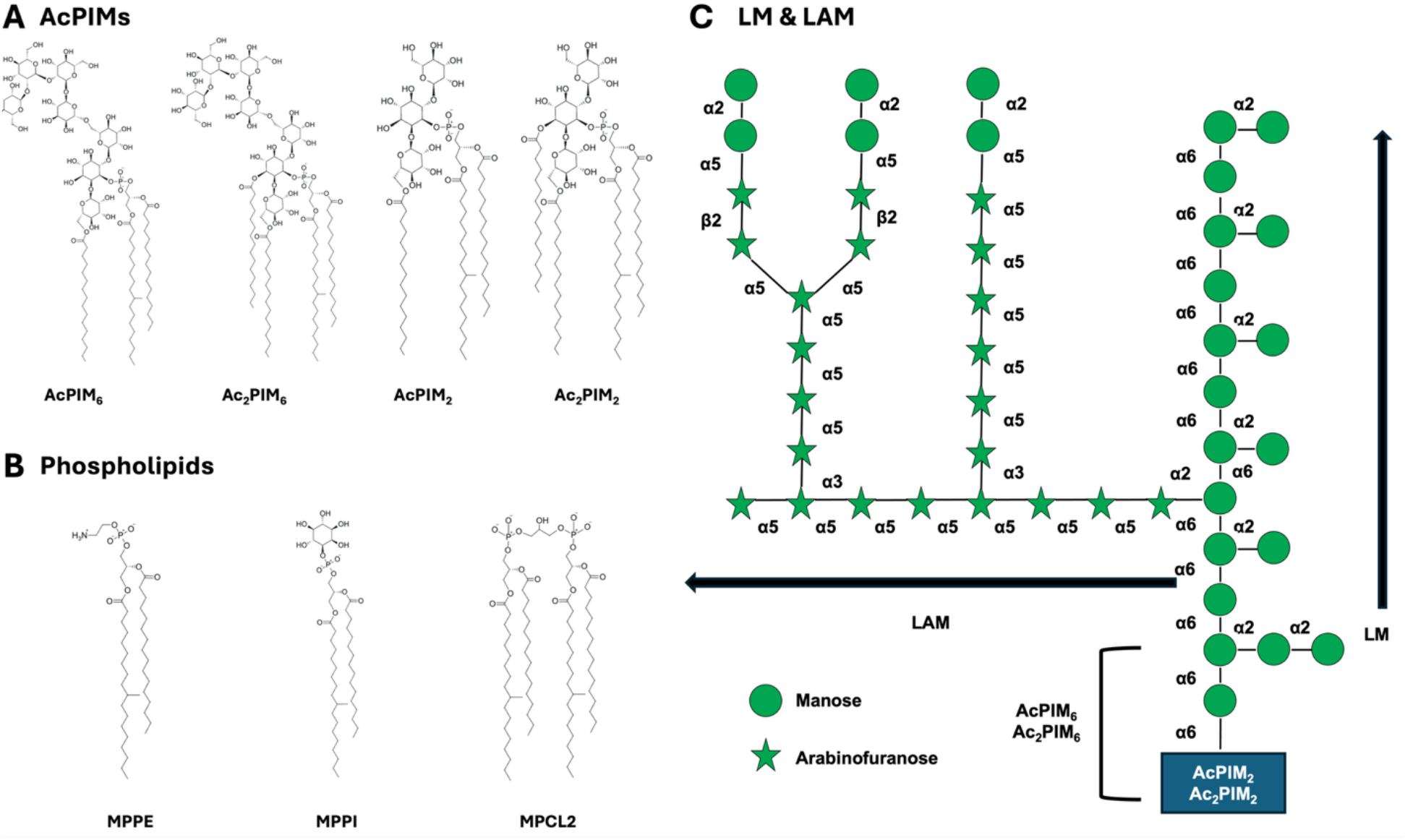
Chemical and symbolic structures of MIM lipids used in this study. (A) Chemical structures of AcPIM_2_, Ac_2_PIM_2_, AcPIM_6_, and Ac_2_PIM_6_ glycolipids, illustrating variations in glycosylation and acyl chain composition. (B) Chemical structures of phosphatidylethanolamine (PE), phosphatidylinositol (PI), and cardiolipin (CL) phospholipids: 10-methyl octadecanoyl (18:0 10-methyl) palmitoyl (16:0) PE (MPPE), MPPI, and MPCL2 with a net charge of –2*e*. (C) LM/LAM lipoglycan carbohydrate sequences.

Several unresolved questions limit our understanding of how LM and LAM function within the MIM. The relative abundance of LM and LAM in the MIM is not precisely known and may vary with species, growth conditions, or environmental stress^22,23^. Moreover, the MIM is a fluid and compositionally heterogeneous environment with enriched PIMs that constitute a substantial fraction of total membrane lipids. Multiacylated PIM species such as Ac_2_PIM_2_, which contains four fatty acyl chains (**Figure 1**), are thought to promote high packing density and reduced membrane fluidity^24–27^, potentially contributing to intrinsic drug resistance by limiting antibiotic influx. Within this crowded and heterogeneous lipid environment, it remains unclear how large, flexible lipoglycans such as LM and LAM behave and how their polysaccharide domains interact with surrounding lipids and with each other. Furthermore, the extent to which LAM reaches above the inner membrane and its interaction with the peptidoglycan and outer membrane is unresolved. While experimental studies have provided valuable information on chemical composition and overall architecture, they offer limited access to molecular-scale structure and dynamics. For example, recent work has revealed pronounced lateral heterogeneity within mycobacterial membranes, including differences in fluidity between cell poles and sidewalls, and disruption of these patterns in LAM-deficient mutants^9,28,29^. However, experimental approaches generally provide ensemble-averaged or static views, making it difficult to resolve molecular-level features such as conformational flexibility of polysaccharide chains, time-dependent clustering, and coupling between glycoconjugates and the underlying lipid leaflets.

Molecular dynamics (MD) simulation provides a means to address these challenges by enabling direct observation of membrane system’s structure and dynamics at the molecular level.

The computational study of complex bacterial membranes has been transformed by advances in both all-atom and coarse-grained MD simulations^9,30–34^. Significant efforts have been devoted to the Gram-negative outer membrane^35–39^, where the development of specialized tools such as the CHARMM-GUI^31,40–42^ LPS Modeler and Membrane Builder has enabled the systematic construction of chemically realistic membrane models. More recent simulations^24,43^ have expanded this framework to include diverse glycolipid chemistries and their interactions with membrane proteins, revealing how variations in sugar headgroups and acylation patterns dictate membrane organization and dynamics.

Extending these computational approaches to the mycobacterial cell envelope presents unique challenges due to the exceptional size, flexibility, and branching complexity of LM and LAM. While the mycobacterial outer membrane (MOM) has begun to receive increased attention through recent all-atom and coarse-grained simulation studies^2,44^, which provide a transformative molecular architecture of the MOM, the MIM, in its full complexity, remains comparatively less explored in simulation at full compositional and atomistic detail. Recent biophysical studies have underscored the importance of lipid domains, PIM enrichment, and membrane heterogeneity in regulating MIM fluidity and host–cell interactions^29,45^, emphasizing that LM and LAM are not merely passive membrane anchors but active regulators of the periplasmic environment. Furthermore, recent computational perspectives have highlighted the necessity of moving from individual lipid models toward complex, multi-component assemblies to accurately capture these emergent properties^2^. Despite these advances, a high-resolution, time-dependent description of how varying LM and LAM ratios influence the physical state of the MIM is still lacking.

In this study, we address these gaps by using all-atom MD simulations to examine various MIM model systems containing different LM/LAM, phospholipid, and PIM ratios to capture both membrane-level properties and sugar-level structural and dynamic behavior. By explicitly incorporating these large, flexible lipoglycans into a realistic environment of phospholipids and PIMs, our approach allows us to capture both global membrane organization and stability and local molecular-scale dynamics that are difficult to access experimentally. By characterizing how these glycoconjugates influence membrane structure and exhibit dynamic behavior within a realistic lipid environment, this study seeks to clarify their roles in mycobacterial membrane organization and to provide a framework for interpreting experimental observations and future MIM modeling and simulation with various (peripheral) membrane proteins.

## Methods

### System Building and Simulation

In this work, we have employed a stepwise modeling strategy to build complex asymmetric MIM systems. Initial simulations focus on symmetric membrane systems of both the MIM inner and outer leaflets. These systems allow us to establish baseline behavior and assess the stability of each inner leaflet symmetric and outer leaflet symmetric model system. We have then extended the simulations to asymmetric membranes using the information from both symmetric systems to better reflect a realistic asymmetric MIM. Comparison between symmetric and asymmetric systems enables us to isolate the effects of leaflet asymmetry on membrane properties and the influence of LM and LAM. **Table S1** summarizes the simulation system information in this study: 3 inner leaflet symmetric systems, 6 outer leaflet symmetric systems, and 9 asymmetric MIM systems. **Figure 1** shows all lipids used in this study: MPPE, MPPI, MPCL2, AcPIM_2_, and Ac_2_PIM_2_ in the inner leaflet and MPPE, MPPI, MPCL2, AcPIM_2_, Ac_2_PIM_2,_ AcPIM_6_, Ac_2_PIM_6_, LM, and LAM in the outer leaflet. All membrane systems were built and equilibrated using the standard CHARMM-GUI *Membrane Builder*^*31,42*^ protocol. We used 3 replicas for each system.

For the inner leaflet symmetric systems, both equilibration and production simulations were carried out in the NPT (constant particle number, pressure, and temperature) ensemble using OpenMM^46^, with the CHARMM36 force field^47^, TIP3P water^48^, and 150 mM KCl. The temperature was maintained at 313.15 K using Langevin dynamics^49^, and the pressure was controlled at 1 bar under semi-isotropic conditions using a Monte Carlo barostat^50^ with a coupling frequency of 5 ps^−1^. All production simulations were run for 500 ns. A 4 fs time step was used with hydrogen mass repartitioning (HMR)^51^, and trajectory frames were saved every 100 ps. Van der Waals interactions were treated with a force-switching scheme between 10 and 12 Å^47^, while long-range electrostatic interactions were calculated using the particle mesh Ewald (PME) method^52^.

To construct the outer leaflet symmetric systems, initial area-per-lipid (APL) values were estimated based on previous coarse-grained simulation studies^9^. Because the exact outer leaflet composition is not known, multiple LM/LAM ratios were tested. In total, six symmetric systems containing LM/LAM were prepared (see **Table S1**). Simulations were started at the lowest LM/LAM concentration and then extended to higher concentrations after careful examination of equilibration behavior. During this process, the ratio of AcPIM_2_/Ac_2_PIM_2,_/AcPIM_6_/Ac_2_PIM_6_ lipids was accordingly adjusted as part of the composition testing. Finally, 9 asymmetric membrane systems were constructed by matching the system membrane area (i.e., the XY) of the three equilibrated inner leaflet symmetric compositions and three representative outer leaflet compositions (see **Table S1**).

Due to the system size, all production simulations of the systems containing LM/LAM were performed using GROMACS^53^ to accommodate larger system sizes in the presence of LM and LAM. To ensure comparability with OpenMM-based simulations of smaller systems, all simulation parameters were kept identical across both programs, including the CHARMM36 force field^47^, TIP3P water model^48^, and 150 mM KCl ionic concentration. The leap-frog integrator was used with a 4 fs time step with the HMR method^51^, and each simulation was run for 1 µs in the NPT ensemble. Electrostatic interactions were treated using the PME method^52^ with a real-space cutoff of 12 Å. Van der Waals interactions were calculated using a cut-off scheme with force-switching applied between 10 and 12 Å. Neighbor lists were generated using the Verlet scheme^54^ and updated every 20 steps with a cutoff of 12 Å. The temperature was maintained at 313.15 K using the Nosé–Hoover thermostat^55,56^ with a coupling constant of 1.0 ps, with separate coupling groups defined for the membrane and solvent. Pressure was controlled semi-isotropically using the Parrinello–Rahman barostat^57^, with a reference pressure of 1 bar, a coupling constant of 5.0 ps, and a compressibility of 4.5 × 10^−5^ bar^−1^ in both the lateral and normal directions. All bonds involving hydrogen atoms were constrained using the LINCS algorithm^58^. Center-of-mass motion was removed every 100 steps using linear correction, applied separately to the membrane and solvent groups. All molecular visualizations and figures were generated using VMD^59^.

### Analysis

To examine the MIM organization along the membrane normal, number density profiles were calculated along the Z-axis. The simulation box was partitioned into discrete slices (bins) of 1 Å width. To ensure accurate alignment throughout the trajectory, the bilayer center was translated to Z = 0 for every frame. The density within each bin at position Z was calculated by counting the number of selected atoms and normalizing it by the volume of the bin. The density in each bin was averaged over the last 500 ns trajectories to produce a time-averaged profile.

Lateral diffusion of lipids was quantified by calculating the mean squared displacement (MSD) from the trajectories using custom Python scripts utilizing the MDAnalysis^60,61^ library. To account for the inherent asymmetry or potential environmental differences between the two leaflets, lipids were first categorized into each leaflet one based on their center-of-mass (COM) position relative to the membrane midplane. To ensure accurate MSD tracking across periodic boundary conditions, the trajectories were unwrapped, effectively removing artificial jumps as lipids crossed the simulation box boundaries. The lateral MSD was then calculated for each lipid type using their XY COM positions:

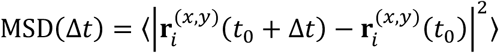

where 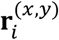 represents the lateral position of lipid *i*at a given time. The MSD was averaged over all lipids within the respective leaflet and over all possible time origins (*t*_0_) to improve statistical convergence. The lateral diffusion coefficient (*D*) was then determined by performing a linear regression on the MSD vs. time plot. In accordance with the Einstein relation for two-dimensional diffusion, the slope of the linear regime was used:

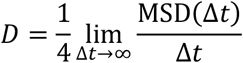

To avoid artifacts at very short timescales or poor sampling at very long timescales, the fit was performed over a robust linear range with the final results reported in cm^2^/s.

To characterize the structural order and fluidity of the lipid bilayers, the deuterium order parameter (S_CD_) was calculated for the hydrocarbon tails using a custom analysis class built on the MDAnalysis framework. For each carbon atom in the acyl chains, the unit vector was defined along the bond connecting the carbon to its attached hydrogen atom. The order parameter was then calculated using following equation:

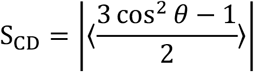

where *θ* is the angle between the C-H bond vector and the bilayer normal, and the angled brackets denote an ensemble average over all equivalent lipids and simulation frames. When multiple hydrogen atoms were bonded to a single carbon, the S_CD_ contribution was averaged over the C-H bonds. S_CD_ values typically range from 0 (indicating complete disorder/isotropy) to 0.5 (indicating perfect alignment perpendicular to the membrane normal). Statistical errors were calculated as the standard deviation of the S_CD_ across the analyzed trajectory. After the calculation, the results of each carbon tails were plotted within each carbon tail and the average of C22 (the first acyl chain carbon bonded to the carbonyl group in the *sn*-2 chain) in each tail was plotted for the comparison. S_CD_ values of each tail (**Figure S3**) were separated, calculated, and plotted initially (**Figure S4**) to choose the proper carbon selection for the comparison.

To identify distinct conformational states of LAM, hierarchical clustering analysis was performed based on pairwise structural similarities. The analysis was implemented using a custom Python workflow incorporating the Bio.PDB module for structural manipulation and the SciPy library for cluster analysis. The similarity between any two conformations was quantified using the root mean square deviation (RMSD). Prior to each comparison, structures were optimally aligned using the Kabsch algorithm to minimize the RMSD and remove differences due to global translation and rotation. A symmetric pairwise distance matrix was constructed, where each entry represents the RMSD between two conformations. After clustering, the final overlap structures were rendered by overlapping clustered structures using VMD.

The hydrophobic thickness of the lipid bilayer was calculated using the MDAnalysis library. To consider the periodic boundary condition, the membrane center of mass was centered at the origin for each frame. The bilayer was partitioned into upper lower leaflets based on the Z-coordinates of the lipid headgroup-tail interface. The hydrophobic thickness was defined as the distance (*d*) between the average Z-position of the C22 and C32 atoms in the top and bottom leaflets. The final values were reported as an average across the analyzed trajectory.

To characterize the LAM conformations and orientations, we calculated LAM tilt angle, residue-residue distance, and lipid contacts. The tilt angle of specific LAM residue relative to the membrane normal (Z-axis) was calculated using CHARMM^62^. The tilt angle was defined by a vector connecting two reference points: the COMs of residues AMAN10 and AARB29 of LAM shown in **Figure S1**. An inter-residue Z-distance was measured through two steps. Initially, Z-coordinates of specific residues (either AMAN10 or AARB29) were extracted using in-house Python script. Then, the distance between two Z-coordinates was calculated by subtracting the two values. To quantify contacts between LAM and the membrane, the average number of terminal sugar contacts per lipid was derived from the final 300 ns. A membrane contact was defined as when any of the sugar residues (AMAN37, AMAN45, and AMAN49 in **Figure S1**) were within 5 Å of a membrane lipid. To determine the volume taken up by LM/LAM above the membrane, a voxel grid composed of 2 × 2 × 2 Å^3^ units was constructed. The van der Waals radius of each atom, with the addition of 1.4 Å representing the radius of water, was used to determine the fraction of occupied voxels in 2 Å slices along the Z axis.

## Results and Discussion

### Mycobacterial inner leaflet symmetric membranes are fluid

To define baseline membrane behavior, we first examined symmetric bilayer models of the MIM inner leaflet. The bilayers consisted of MPPE, MPPI, MPCL2, AcPIM_2_, and Ac_2_PIM_2_, allowing us to focus on the glycolipid composition expected to dominate the inner leaflet^9,27,63,64^. We varied the relative amount of AcPIM_2_ and Ac_2_PIM_2_ across three compositions (I-1:1, I-1:2, and I-2:1; see **Table S1** for the system name and the corresponding lipid composition) to examine how changes in AcPIM_2_ and Ac_2_PIM_2_ contents affect lipid packing and dynamics in the absence of larger glycoconjugates. Thus, these symmetric bilayers establish a simple reference system that can be used to evaluate additional effects of LM/LAM and membrane asymmetry in later simulations.

All three membrane compositions form intact bilayers and remain stable with no pore formation, large-scale deformation, or lateral phase separation (**Figure 2A**). The density profiles along the membrane normal (i.e., the *Z*-axis) show a clear hydrophobic core in all systems with the headgroup regions at ±22 Å and similar bilayer hydrophobic thicknesses across compositions (**Figure S2**). The headgroup regions are crowded with phosphates, glycerols, and sugar rings, while the lipid tails from both leaflets meet at the membrane midplane (Z = 0), creating a dip. The density profiles also clearly reflect the order in the number of acyl chains of each lipid type: MPCL2 and Ac_2_PIM_2_ with 4 tails > AcPIM_2_ with 3 tails > MPPE and MPPI with 2 tails (see **Figure 1**).

**Figure 2.**
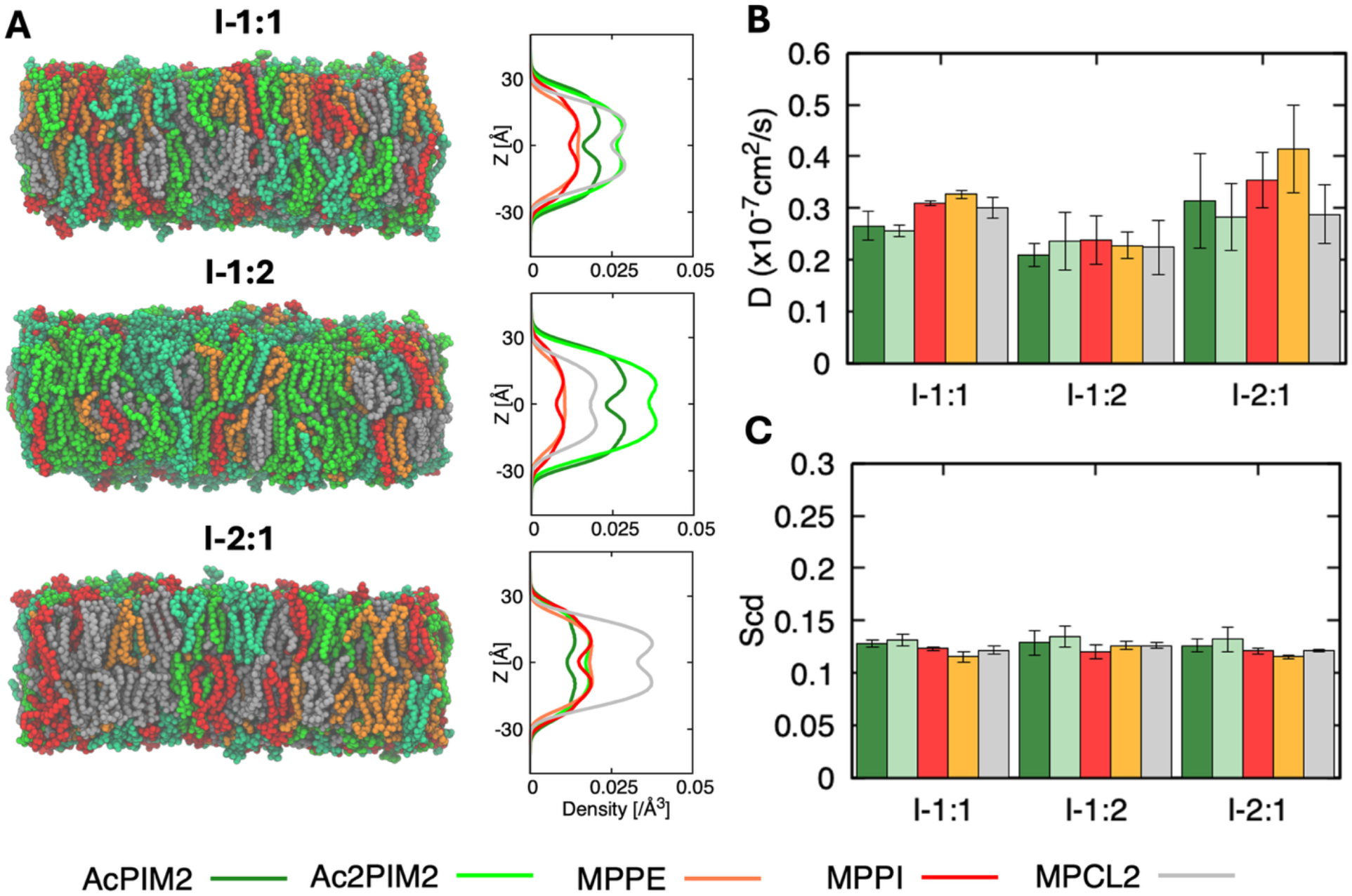
Inner leaflet symmetric membrane structure and dynamics. (A) Representative equilibrated snapshots and corresponding number density profiles along the membrane normal (i.e., the Z-axis) for system I-1:1, I-1:2, and I-2:1 (see **Table S1** for the system name and the corresponding lipid composition) composed of phospholipids (MPPE, MPPI, MPCL2) and phosphatidylinositol mannosides (AcPIM_2_ and Ac_2_PIM_2_). Each lipid is depicted (B) Lateral diffusion coefficients of individual lipid species obtained from mean squared displacement analysis. (C) Deuterium order parameters (S_CD_) of C22 carbon of phospholipid acyl chains. All systems remain structurally stable and fluid across the compositions studied. Error bars represent standard deviations among 3 replicas.

The lateral diffusion coefficients for all lipid species fall in the range of ∼2-4 × 10^-8^ cm^2^ s^-1^, consistent with lipids in fluid phase membranes (**Figure 2B**), which is typically reported to be on the order of 10^-8^ ∼ 10^-7^ cm^2^s^-1^ in both model and biological lipid bilayers^30,65^. While a slight increase in the mean diffusion coefficients for MPPI and MPPE is observed in the I-2:1 system, these variations remain within the calculated error margins, indicating that the overall lateral mobility remains statistically comparable across the different membrane compositions in this study. Increasing PIM content leads to a moderate decrease in diffusion, likely due to increased steric constraints at the membrane surface. Within each system, however, phospholipids and PIMs show similar mobilities, and acyl chain differences between AcPIM_2_ and Ac_2_PIM_2_ do not noticeably affect diffusion. Consistent with this, deuterium order parameters (S_CD_) are nearly identical across all composition, indicating that changes in PIM abundance do not significantly alter acyl chain packing (**Figure 2C**). The stability of these values suggests that the slight variations observed in lateral diffusion (**Figure 2B**) are likely governed by surface-level headgroup interactions or long-range lateral spacing rather than a fundamental change in the hydrophobic core’s ordering. Overall, these results show that the phospholipid-PIM membrane remains dynamically fluid and structurally stable over the compositions studied, providing a baseline for subsequent simulations including LM and LAM.

### Progressive enrichment of LM and LAM in outer leaflet symmetric membranes systematically reduces lipid mobility

Building on the symmetric inner leaflet models, we next examined symmetric outer leaflet membranes containing the larger glycoconjugates LM and LAM (**Figure 3A**). As the exact lipid ratio of the MIM outer leaflet is unknown, we constructed 6 model systems with increasing LM/LAM concentration (see **Table S1** for the system name and the corresponding lipid composition). All outer leaflet symmetric membrane systems remain stable over microsecond simulation timescales, but clear composition-dependent differences in structure and dynamics are observed. In each system, phospholipids (MPPE, MPPI, and MPCL2) and PIMs form a well-defined hydrophobic core with similar membrane thicknesses (**Figure S2**), while LM and LAM glycans occupy an extended interfacial region. As their content increases from O-10:2 to O-10:10, the density profiles along the membrane normal show a gradual broadening of the outer leaflet distribution.

**Figure 3.**
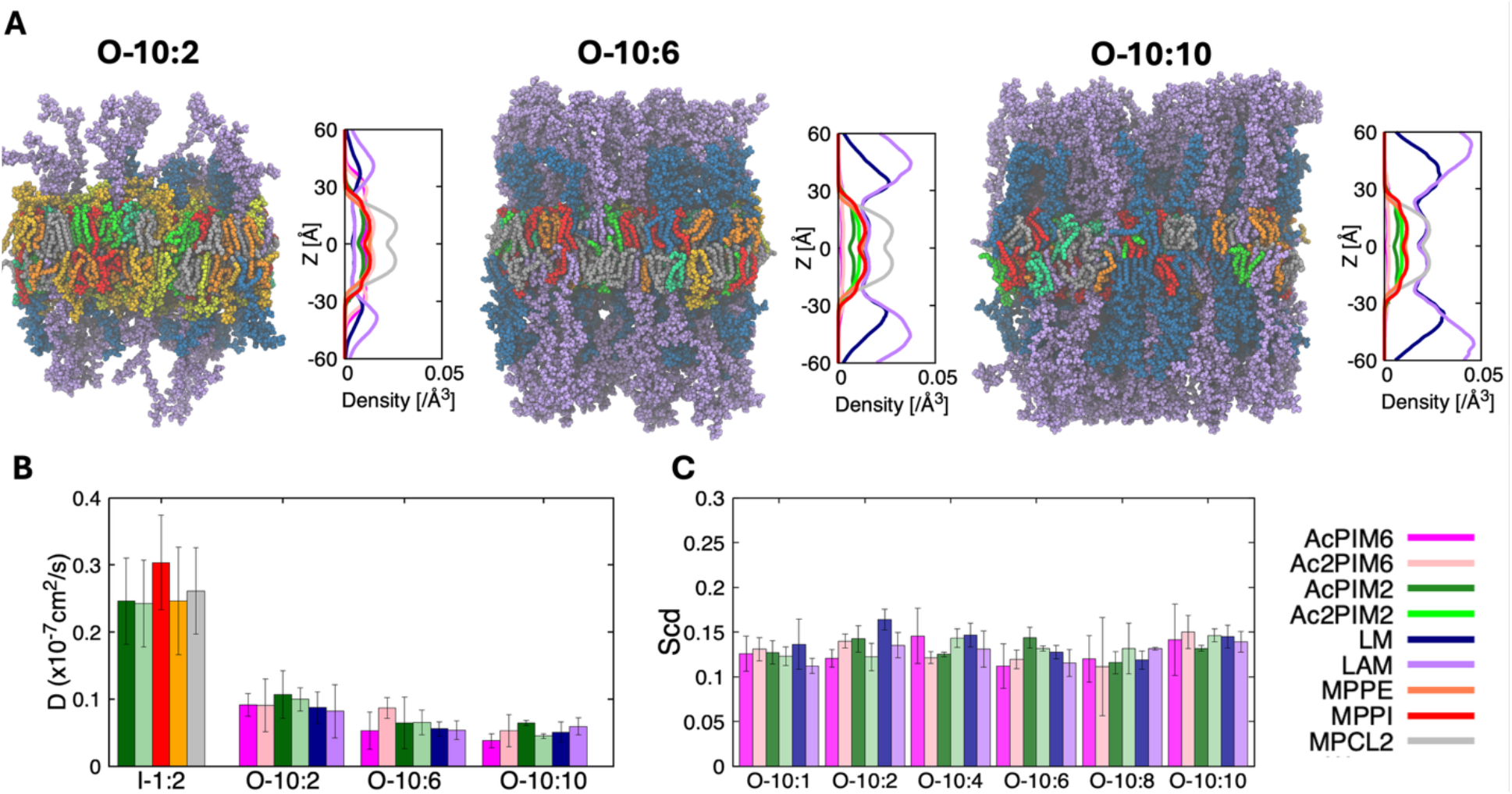
Structure and dynamics of outer leaflet symmetric membranes containing LM and LAM. (A) Representative equilibrated snapshots and number density profiles along the membrane normal for the outer leaflet symmetric membrane systems with increasing LM/LAM content (see **Table S1** for the system name and the corresponding lipid composition). Each lipid is depicted with separated colors: orange for MPPE, red for MPPI, gray for MPCL2, dark green for AcPIM_2_, light green for Ac_2_PIM_2_, magenta for AcPIM_6_, yellow for Ac_2_PIM_6_, dark blue for LM, and purple for LAM. (B) Lateral diffusion coefficients of individual lipid species for the inner-leaflet reference system (I-1:2) and outer leaflet symmetric membranes (O-10:2, O-10:6, and O-10:10). (C) Deuterium order parameters (S_CD_) of C22 carbon of phospholipid acyl chains as a function of LM/LAM content. Error bars represent standard deviations among 3 replicas.

LM and LAM organize into an extended polysaccharide layer that is solvated, as evidenced by the significant exposure of carbohydrate above the membrane hydrophobic area (**Figure 3A** and **Figure 5A**). Both density analysis and visual inspection indicate that LM glycan chains adopt relatively compact but flexible conformations, whereas LAM forms larger and more diffuse structures due to its longer, branched arabinan domains. In systems with higher LM/LAM content (O-10:6 and O-10:10), neighboring carbohydrate chains interact with each other substantially, giving rise to a crowded, brush-like layer above the membrane surface. Despite this pronounced surface restructuring, the lipid acyl chains remain well packed, indicating that incorporation of LM and LAM does not destabilize the bilayer core. Instead, their primary effects are confined to the membrane surface and periplasmic space.

Lipid mobility in the outer leaflet is strongly dependent on both the concentration and identity of the glycoconjugates. As shown in **Figure 3B**, the lateral diffusion coefficients decrease monotonically with increasing LM/LAM content for all lipid species, including phospholipids and AcPIMs. The reduction in diffusion is most pronounced in LAM-rich systems, consistent with the larger size and higher branching of LAM relative to LM. These trends suggest that glycoconjugates restrict lateral motion through steric crowding between glycan chains, resulting in a less dynamic outer leaflet at high glycoconjugate loadings. On the other hand, lipid packing shows a very similar result, regardless of the LM/LAM content (**Figure 2C** and **Figure 3C**). Together, these results point to the development of lateral heterogeneity with glycoconjugate-enriched regions exhibiting higher rigidity and reduced dynamics. Overall, LM and LAM act as key regulators of outer leaflet structure and dynamics, producing a thick, hydrated surface layer that likely contributes to reduced permeability, enhanced mechanical resilience, and modulation of host-pathogen interactions.

### Molecular crowding reshapes the conformational and orientational landscape of LAM

The metrics discussed so far establish the general stability of both MIM inner leaflet and outer leaflet symmetric membrane systems. We expect that the LAM surface density does not just alter global properties, but it restricts the polysaccharide domains movement, orientation, and the solvent-accessible volume in the periplasmic space due to physical crowding. **Figure 4A** shows representative snapshots and structural overlays of LAM from the low and high concentration systems. At low concentration (O-10:1), LAM molecules are highly flexible and extended parallel to the membrane surface, and the sugar residues sample a wide range of orientations above the membrane. As its concentration increases to O-10:10, however, these structures become much more compact with less flexibility and align parallel to the membrane normal. The structural overlays indicate that higher LAM surface density physically limits the polysaccharide chains, a clear signature of crowding due to LAM-LAM interactions.

**Figure 4.**
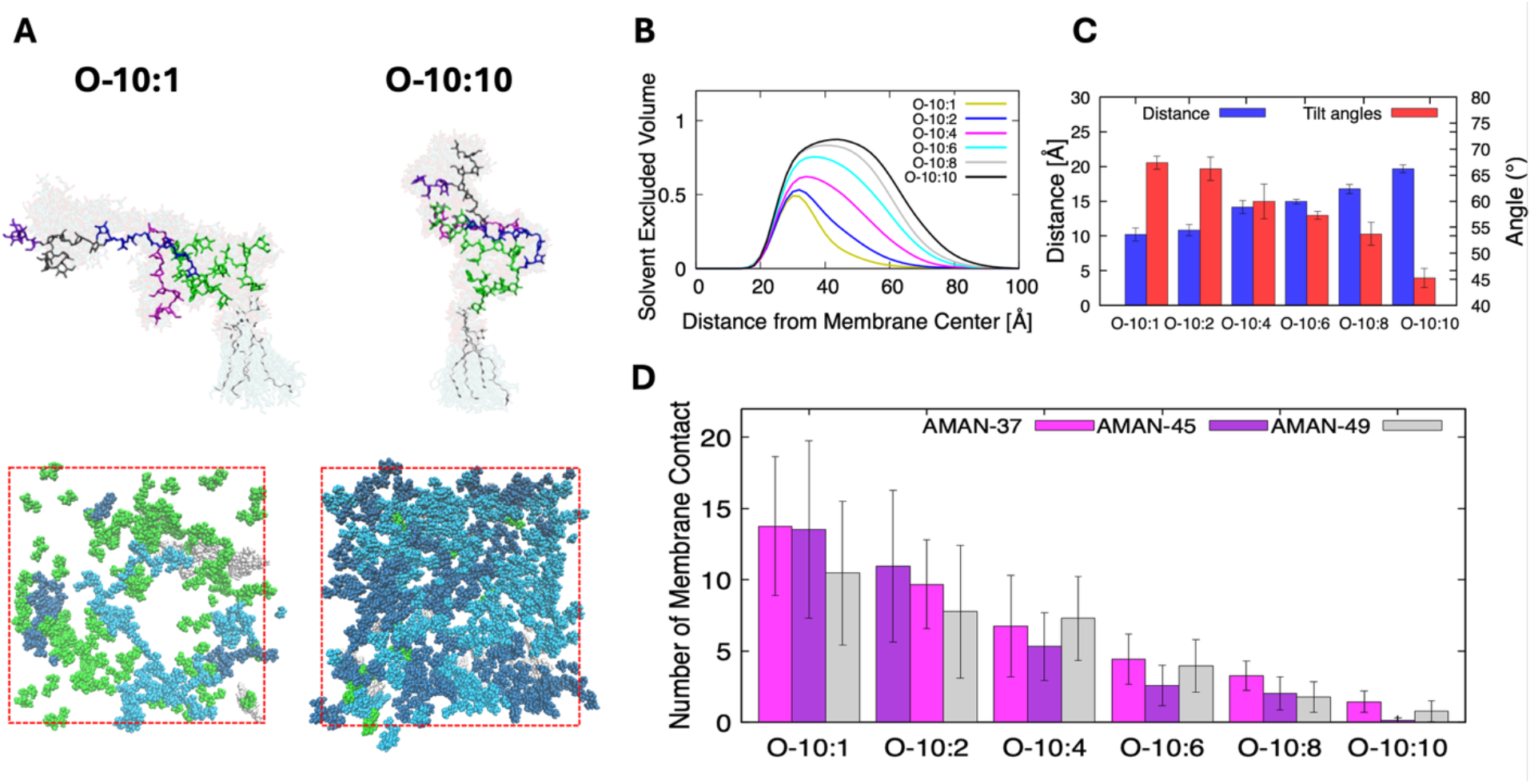
Concentration-dependent structure and dynamics of LAM in a mycobacterial inner membrane. (A) Representative snapshots and overlaid conformations of LAM extracted from low and high LAM concentration systems. Red dashed boxes indicate the primary simulation system XY area. (B) Solvent excluded, LM/LAM-occupied volume profiles as a function of distance from the membrane center (Z = 0) for systems O-10:1, O-10:2, O-10:4, O-10:6, O-10:8, and O-10:10. A value of 1 represents that all volume is occupied by LM/LAM and thus inaccessible to solvent (C) Average Z distance between LAM sugar residues AMAN10 and AARB29 in **Figure S1** (blue bars, left axis) and average tilt angle of a vector from AMAN10 to AARB29 relative to the membrane normal (red bars, right axis) for systems O-10:1 to O-10:10. (D) Average number of contacts formed between LAM sugar residues (defined in **Figure S1**) and the membrane lipids for systems O-10:1 to O-10:10. Error bars represent standard deviations among 3 replicas.

**Figure 5.**
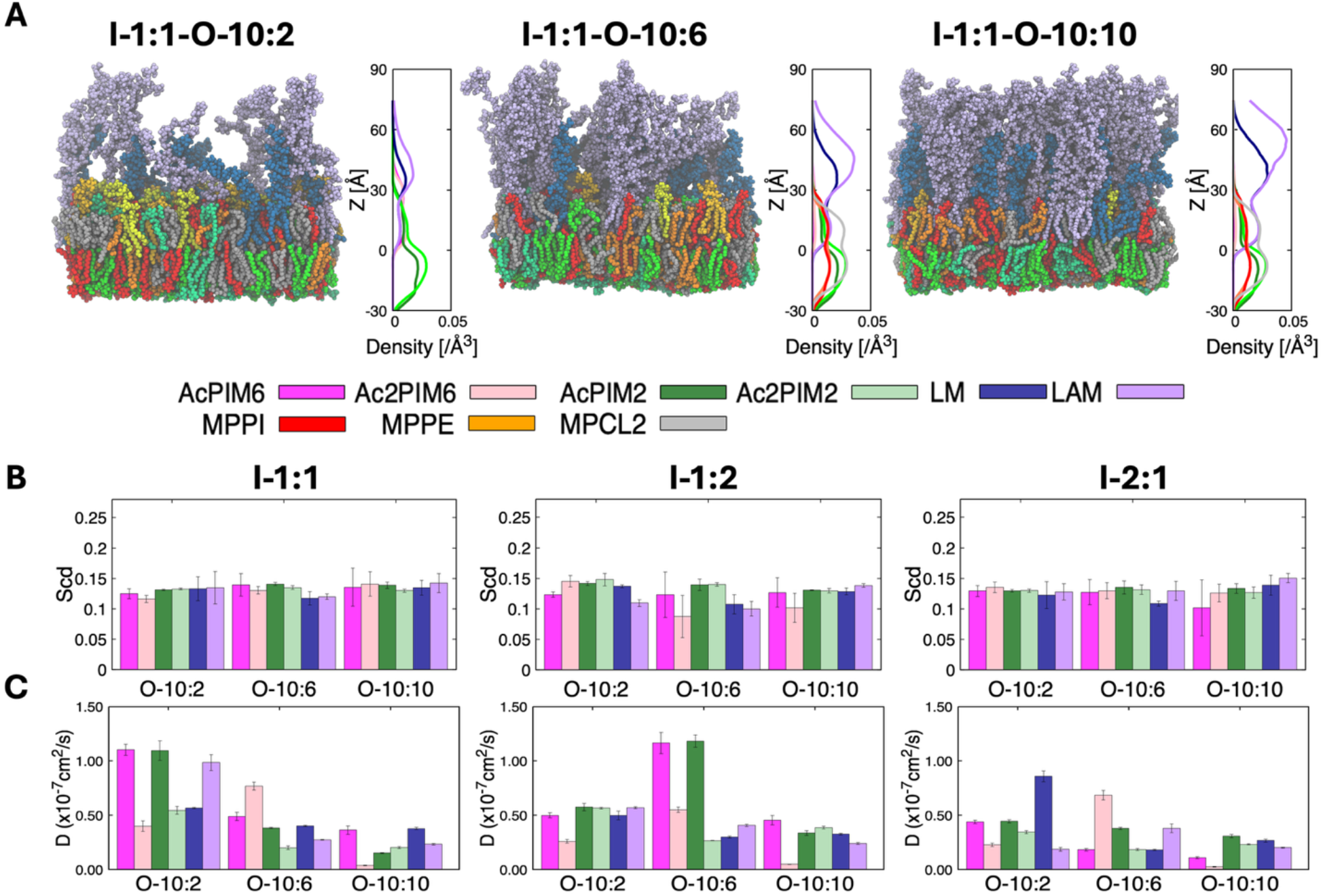
Structure and dynamics of asymmetric mycobacterial inner membrane systems. (A) Representative snapshots and number density profiles illustrating leaflet asymmetry. (B) Lipid acyl chain order parameters (S_CD_) of C22 carbon of acyl chains for outer leaflet phospholipids. (C) Lateral diffusion coefficients of lipid species in asymmetric bilayers. Increasing outer leaflet lipoglycan content reduces lipid mobility in both leaflets, demonstrating long-range dynamic coupling mediated by the lipoglycan-rich outer leaflet.

As LAM concentration increases, steric crowding above the membrane forces the glycan chains to become more vertical and reach further into the periplasmic space. This elongation is visible in the average Z distance between the sugar residues AMAN10 and AARB29 (**Figure 4C**). As we move from O-10:1 to O-10:10, the mean Z distance increases, demonstrating that the terminal sugars are moving farther from the membrane. This is simultaneously accompanied by a decrease in the average tilt angle of a vector from AMAN10 to AARB29 (**Figure 4C**). Essentially, as LAM concentration increases, the polysaccharide chains reorient themselves to lie more perpendicular to the membrane surface. We calculated the solvent excluded volume (SEV), i.e., LM/LAM-occupied volume relative to the distance from the membrane midplane to quantify LM/LAM’s spatial occupancy (**Figure 4B**). In the low concentration systems, the SEV distributions are low, confirming that the sugar residues are largely solvent exposed, free to move, and stay primarily near the membrane-solvent interface. In the O-10:8 and O-10:10 systems, however, the SEV rises significantly, especially further away from the membrane. This agrees with the observed elongation of LAM as concentration increases. At these higher concentrations, LM/LAM takes up a significant portion of space above the membrane (∼80%), providing a significant barrier to molecular permeability.

Finally, we examine the number of atom contacts between LAM sugars and the lipid molecules (**Figure 4D**). These interactions are biologically relevant given that arabinosyltransferases involved in LAM biosynthesis are membrane-associated enzymes, implying that transient contacts between LAM carbohydrate domains and the membrane surface is necessary during LAM elongation^66–68^. Our analysis shows that the O-10:1 and O-10:2 systems have the highest number of contacts. This is expected as LAM at low concentration has the most conformational freedom to bend down to make membrane contact. As concentration increases, the number of contacts per molecule drops significantly. At higher LAM surface densities, neighboring LAM molecules crowd one another, reducing the available space for individual carbohydrate chains to spread across the membrane surface. Instead, the chains tend to adopt more upright conformations, which limits their ability to maintain extended contacts with lipids. Under these conditions, interactions between neighboring carbohydrate domains become more prevalent than direct carbohydrate-lipid contacts.

Taken together, these simulations indicate that LAM concentration strongly influences the conformational organization and membrane interaction behavior. At low surface densities, LAM remains flexible, highly solvent-exposed, and capable of frequent membrane contact, whereas crowding at higher densities drives a transition toward a more compact, membrane-normal orientation with reduced lipid interactions. Given that LAM abundance fluctuates with physiological state, particularly during the transition to stationary phase, our results suggest that such variations act as a structural switch for cell envelope remodeling. By modulating surface density, the bacteria can effectively control LAM orientation and its subsequent interactions with the host environment^69^. Although LAM and LM are key components of the mycobacterial cell envelope and important virulence factors, their precise spatial organization relative to the arabinogalactan-peptidoglycan complex, and whether LAM extends fully through the peptidoglycan layer in vivo, warrant further investigations^70^.

### Lipid diffusion is primarily dictated by outer leaflet LM and LAM concentration, demonstrating asymmetrical contributions of each leaflet

To extend the symmetric membrane analysis toward a more physiologically relevant model, we constructed asymmetric bilayers in which AcPIM_6_, Ac_2_PIM_6_, LM, and LAM are present exclusively in the outer leaflet (**Figure 5**). As shown in **Table S1**, in our 9 asymmetric MIM models, the inner leaflet is composed of phospholipids, AcPIM_2_, and Ac_2_PIM_2_ at the same ratios used in the symmetric inner leaflet models, while the outer leaflet has increasing levels of LM and LAM corresponding to O-10:2, O-10:6, and O-10:10 compositions. Given the limited experimental data on MIM lipid composition, particularly the relative ratios of glycoconjugates (AcPIMs, LM, and LAM), these systems also serve to evaluate a range of reasonable inner-outer leaflet composition combinations. This design allows direct assessment of leaflet coupling in the asymmetric MIMs. All asymmetric MIM systems remain stable over microsecond simulations, forming intact bilayers without pore formation or large-scale deformation.

Structure analysis reveals that lipid distribution largely follows trends shown in their respective symmetric systems. The inner leaflet retains a compact hydrophobic core that closely resembles the symmetric inner leaflet systems and shows little sensitivity to changes in the opposing leaflet. The outer leaflet also demonstrates similar trends to the symmetric systems. As the LM and LAM content increases, the Z-dependent density profiles show a gradual extension of the outer leaflet region, consistent with the formation of an increasingly prominent, polysaccharide layer that extends into the solvent (**Figure 5A**). As a result, there is an increase in effective membrane thickness driven primarily by LAM extension away from the membrane surface. LM and LAM carbohydrate domains organize into an extended surface layer whose structure depends on glycoconjugate content. At lower concentrations, individual LM and LAM molecules remain relatively separated, whereas higher one leads to significant contacts between neighboring carbohydrate chains. In LM/LAM-rich systems, this contact produces a dense, brush-like layer at the membrane surface. Despite this pronounced surface crowding, the inner leaflet remains well packed, indicating that bulky outer leaflet glycoconjugates do not disrupt inner leaflet organization or compromise bilayer integrity. Similar acyl chain orders are consistent with these trends (**Figure 5B**; see also **Figure 2C** and **Figure 3C**). For example, outer leaflet AcPIMs stays within a similar range without noticeable differences in order with increasing LM/LAM concentration, maintaining almost identical order parameters.

Lipid dynamics reflects this leaflet-specific behavior, though the lateral diffusion coefficients (*D*) exhibit a more complex dependence on membrane composition than the structural metrics alone. Rather than a linear decrease in mobility, the asymmetric models maintain relatively similar diffusion rates between the O-10:2 and O-10:6 systems. In fact, for certain lipid species, a slight increase in *D* is observed as glycan concentration shifts from O-10:2 to O-10:6 (**Figure 5C**; see also **Figure 3B**). This suggests that at intermediate glycan densities, the membrane environment remains sufficiently fluid to accommodate additional glycans without imposing immediate kinetic restrictions. However, once the surface density is further increased to O-10:10, diffusion coefficients drop significantly across all species, marking a clear transition to a crowding-induced regime where molecular packing significantly limits lateral mobility. In contrast, lipid diffusion does not change while varying the inner leaflet composition. This demonstrates that the outer leaflet, and LM/LAM concentration, has more influence on diffusion than the inner leaflet. (**Figure 5C**; see also **Figure 2B**). This difference highlights a dynamic separation between the two leaflets where LM and LAM play the largest role in dictating lipid diffusion in the outer leaflet.

The observed lateral mobility of LM and LAM molecules presents a distinct physical profile compared to lipopolysaccharides (LPS) found in the outer membrane (OM) of Gram-negative bacteria. All-atom MD simulations of LPS-containing systems have characterized the OM outer leaflet as a nearly immobile, gel-like lattice^71,72^. In such systems, divalent ions such as Mg^2+^ or Ca^2+^ act as an electrostatic glue between the phosphate groups of lipid A, imposing a rigid barrier that severely restricts later diffusion^73,74^. This rigidity is a direct consequence of the divalent cation-mediated cross-linking between LPS phosphate groups. In contrast, the mycobacterial lipoglycans (LM and LAM) lack such strong, charge-driven intermolecular interactions, as the primary interactions between neighboring chains are mediated by transient hydrogen bonding and van der Waals forces^63^. Consequently, LM and LAM remain significantly more mobile than LPS, allowing the mycobacterial inner membrane to retain its fluid character even under high glycoconjugate concentrations. This suggests that the mycobacterial cell envelope achieves its protective barrier properties through a thick, hydrated polysaccharide buffer that maintains a degree of fluid character, potentially facilitating the lateral reorganization of membrane proteins and cell wall biosynthetic machinery. These leaflet-specific dynamics likely serve a dual evolutionary purpose, providing a robust permeability barrier against exogenous antibiotics while preserving the membrane plasticity necessary for host-pathogen interactions and the constant remodeling of the mycobacterial cell wall.

## Conclusions

In this work, we present an all-atom MD simulation study of the MIM with a focus on the role of its major glycolipids (AcPIM_2_, Ac_2_PIM_2,_ AcPIM_6_ and Ac_2_PIM_6_) and lipoglycans (LM and LAM). By systematically varying the concentration of these molecules in both symmetric and asymmetric membrane models, we explore how changes in their loading influence membrane structure, dynamics, and interfacial organization at the molecular resolution.

Experimental studies have long suggested that the MIM (or mycobacterial plasma membrane) differs fundamentally from canonical bacterial plasma membrane, particularly due to the abundance of PIM lipids and large lipoglycans^27,63^. Consistent with this view, our simulations show that the inner leaflet, enriched in phospholipids and PIMs (AcPIM_2_ and Ac_2_PIM_2_), remains in a liquid-disordered state across all conditions examined in this study. Variations in PIM content primarily affect interfacial interactions and lipid organizations, with no significant changes in hydrophobic core packing or bilayer thickness^2^. This behavior aligns with experimental observations reporting preserved membrane fluidity in mycobacteria despite substantial remodeling of lipid composition^29^.

In contrast, the properties of the outer leaflet are strongly modulated by LM and LAM concentration. Increasing lipoglycan abundance leads to reduced lateral mobility and an extension of LAM into the periplasmic space. These findings are consistent with biophysical studies suggesting that LM and LAM form an extended, hydrated layer at the membrane surface and contribute to the unusual thickness and heterogeneity of the mycobacterial envelope^45^. Our results provide a molecular interpretation of these observations, showing how increased surface density of large polysaccharide molecules directly alters membrane dynamics and interfacial structure. At higher surface densities, LAM undergoes a gradual shift from flexible, solvent-exposed conformations to more compact and orientationally ordered states. This crowding-induced reorganization reduces the frequency of direct sugar-lipid contacts while promoting a more uniform alignment of polysaccharide chains with respect to the membrane. Such behavior is consistent with polymer brush models that have been proposed to describe cell-surface glycans and supports the idea that LAM can act as a dynamic, concentration-dependent regulator of the periplasmic-facing environment rather than as a static structural component^75,76^.

These observations reinforce the view that the MIM is an active and highly organized membrane rather than a simple permeability barrier. Previous work has suggested that glycolipid and lipoglycan composition influences antibiotic susceptibility and membrane-associated processes in mycobacteria^15,77^. Our results provide a physical basis for these effects by showing how LAM crowding can limit access to the membrane surface and reshape the interfacial landscape. The coexistence of a fluid inner leaflet and a densely decorated outer leaflet likely enables mycobacteria to balance membrane flexibility with protection under diverse environmental conditions.

Most computational efforts to date have focused on the mycobacterial outer membrane, particularly its mycolic-acid rich architecture and low permeability. By comparison, the inner membrane has received far less attention at comparable resolution. Our results highlight an important distinction between the two membranes. Unlike the rigid outer membrane, the MIM remains fluid even at elevated lipoglycan concentrations^29^. At the same time, both membranes rely on complex carbohydrate parts to modulate organization and dynamics, suggesting a conserved envelope-level strategy in which sugars play a central regulatory role.

Overall, this study provides a molecular framework for understanding how LM and LAM contribute to the physical properties of the mycobacterial inner membrane and complements existing experimental and modeling efforts. The models presented here should aid in interpreting observations related to membrane heterogeneity, altered glycolipid composition, and LAM-deficient phenotypes^15,78^. Future work could extend this approach to include membrane proteins or to explore the effects of environmental variables, such as pH and ionic strength, on glycoconjugate organization. Together, these efforts will help clarify how the unique architecture of the MIM contributes to virulence and drug resistance.

## Acknowledgements

This work is supported by the CNRS-MITI grant “Modélisation du vivant” 2020 (to M.C.), NSF MCB-2111728, and NIH R35 GM153458 (to W.I.).

## Data Availability

The input and restart files necessary for the continuation of all simulations are freely available in a Zenodo dataset (https://doi.org/10.5281/zenodo.19239629).

