## Supporting Information for "Influence of Lipomannan and Lipoarabinomannan Concentration on Mycobacterial Inner Membranes Characterized by All-atom Simulations"

**Table S1.** System composition and simulation details for the symmetric and asymmetric mycobacterial membrane models in this study.[1]

| | Name | Abrev. | MPPE | MPPI | MPCL2 | AcPIM$_6$ | Ac$_2$PIM$_6$ | AcPIM$_2$ | Ac$_2$PIM$_2$ | LM | LAM | CLA | POT | Water | Total Atoms | Initial Size | Time (μs) |
|---|---|---|---|---|---|---|---|---|---|---|---|---|---|---|---|---|---|
| Inner | I-1:2:0:0 | I-1:2 | 80 | 80 | 80 | - | - | 160 | 160 | - | - | 142 | 702 | 50,440 | 275,364 | 163 x 163 x 115 | 0.5 |
| | I-1:1:0:0 | I-1:1 | 112 | 112 | 112 | - | - | 112 | 112 | - | - | 138 | 698 | 49,436 | 264,616 | 161 x 161 x 115 | |
| | I-2:1:0:0 | I-2:1 | 140 | 140 | 140 | - | - | 70 | 70 | - | - | 135 | 695 | 48,341 | 254,563 | 158 x 158 x 115 | |
| Outer | O-10:5:8:1 | O-10:1 | 80 | 80 | 80 | 64 | 64 | 40 | 40 | 8 | 8 | 493 | 957 | 172,779 | 640,403 | 150 x 150 x 311 | 1 |
| | O-10:5:6:2 | O-10:2 | 80 | 80 | 80 | 48 | 48 | 40 | 40 | 16 | 16 | 444 | 892 | 156,680 | 596,328 | 150 x 150 x 295 | |
| | O-10:4:2:4 | O-10:4 | 80 | 80 | 80 | 16 | 16 | 40 | 40 | 32 | 32 | 445 | 861 | 156,237 | 603,641 | 148 x 148 x 308 | |
| | O-10:4:1:6 | O-10:6 | 80 | 80 | 80 | 8 | 8 | 32 | 32 | 48 | 48 | 439 | 855 | 154,654 | 619,728 | 153 x 153 x 301 | |
| | O-10:4:0.5:8 | O-10:8 | 80 | 80 | 80 | 4 | 4 | 32 | 32 | 64 | 64 | 465 | 913 | 165,385 | 682,341 | 163 x 163 x 296 | |
| | O-10:4:1:10 | O-10:10 | 80 | 80 | 80 | 8 | 8 | 32 | 32 | 80 | 80 | 530 | 1010 | 188,979 | 783,621 | 172 x 172 x 306 | |
| Asym. | I-1:2-O-10:2 | | 75 | 75 | 75 | 20 | 20 | 90 | 90 | 10 | 10 | 297 | 762 | 105,062 | 433,705 | 151 x 151 x 210 | 1 |
| | I-1:2-O-10:6 | | 75 | 75 | 75 | 4 | 4 | 86 | 86 | 24 | 24 | 285 | 738 | 101,922 | 437,925 | 153 x 153 x 211 | |
| | I-1:2-O-10:10 | | 75 | 75 | 75 | 5 | 5 | 86 | 86 | 40 | 40 | 293 | 778 | 106,963 | 483,432 | 162 x 162 x 209 | |
| | I-1:1-O-10:2 | | 89 | 89 | 89 | 20 | 20 | 69 | 69 | 10 | 10 | 289 | 754 | 103,042 | 424,248 | 150 x 150 x 209 | |
| | I-1:1-O-10:6 | | 89 | 89 | 89 | 4 | 4 | 65 | 65 | 24 | 24 | 277 | 730 | 99,361 | 426,845 | 152 x 152 x 209 | |
| | I-1:1-O-10:10 | | 89 | 89 | 89 | 4 | 4 | 65 | 65 | 40 | 40 | 288 | 773 | 104,080 | 471,392 | 162 x 162 x 208 | |
| | I-2:1-O-10:2 | | 101 | 101 | 101 | 20 | 20 | 51 | 51 | 10 | 10 | 288 | 753 | 102,198 | 418,816 | 149 x 149 x 211 | |
| | I-2:1-O-10:6 | | 101 | 101 | 101 | 4 | 4 | 47 | 47 | 24 | 24 | 276 | 729 | 98,822 | 422,328 | 150 x 150 x 211 | |
| | I-2:1-O-10:10 | | 101 | 101 | 101 | 4 | 4 | 47 | 47 | 40 | 40 | 286 | 771 | 104,304 | 469,162 | 160 x 160 x 210 | |

[1]All membrane systems investigated in this study include the symmetric inner leaflet models, the symmetric outer leaflet models with varying LM/LAM ratios, and the asymmetric bilayers combining the fixed inner leaflet compositions with the systematically varied outer leaflet lipoglycan content. The number of each lipid species, counterions, total atoms, and production simulation times are reported for each system.

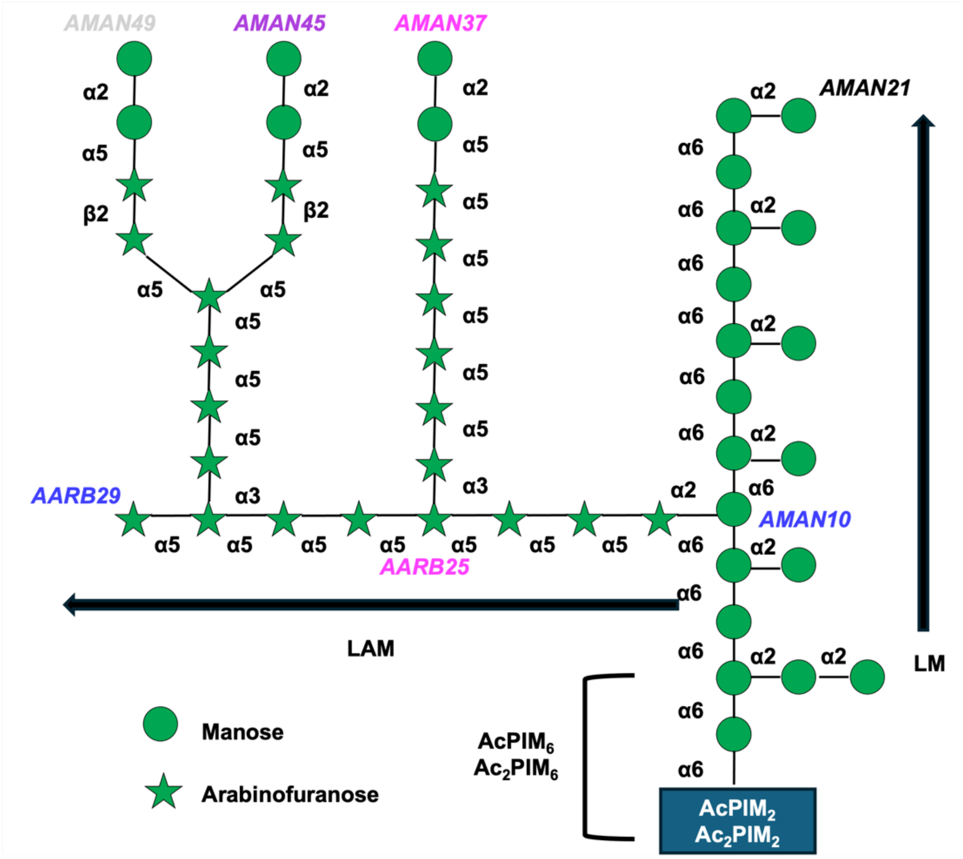


**Figure S1**. LM/LAM residues used for the tilt angle (AMAN10 and AARB29), averaged distance (AMAN10 and AARB29), and membrane contact (AMAN37, AMAN45, and AMAN49) calculations.

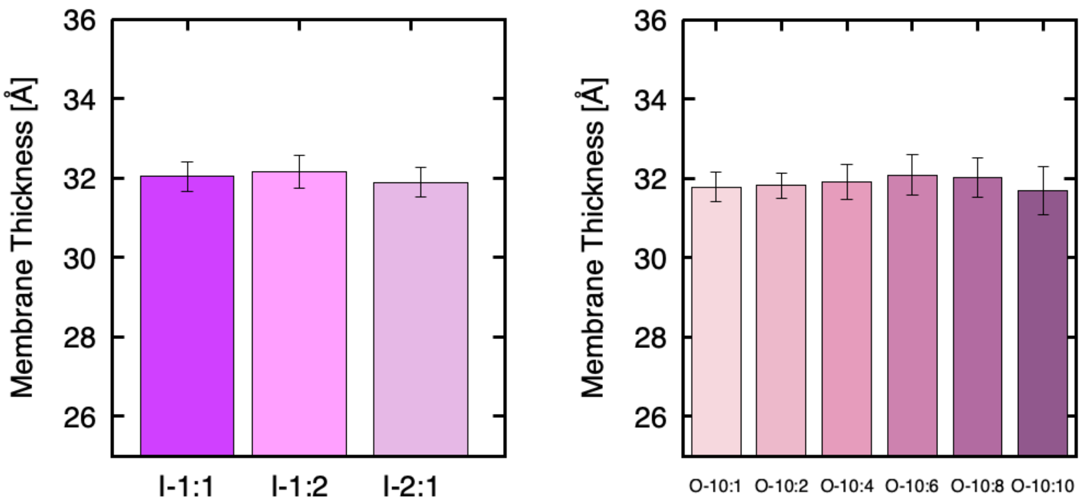


**Figure S2**. Membrane hydrophobic thicknesses of all symmetric systems.

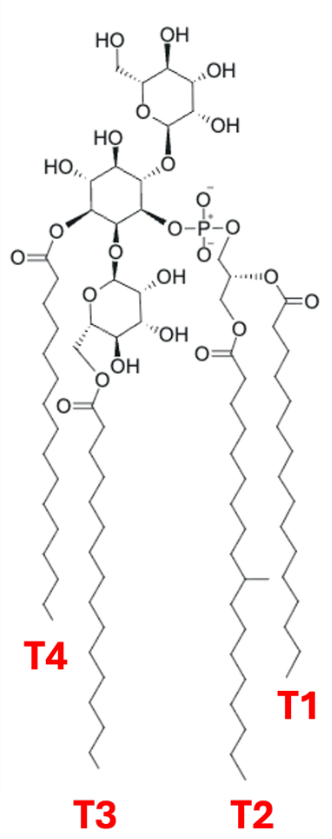


$Ac_2PIM_2$

**Figure S3.** Individual chains used for order parameter calculations of LM and LAM.

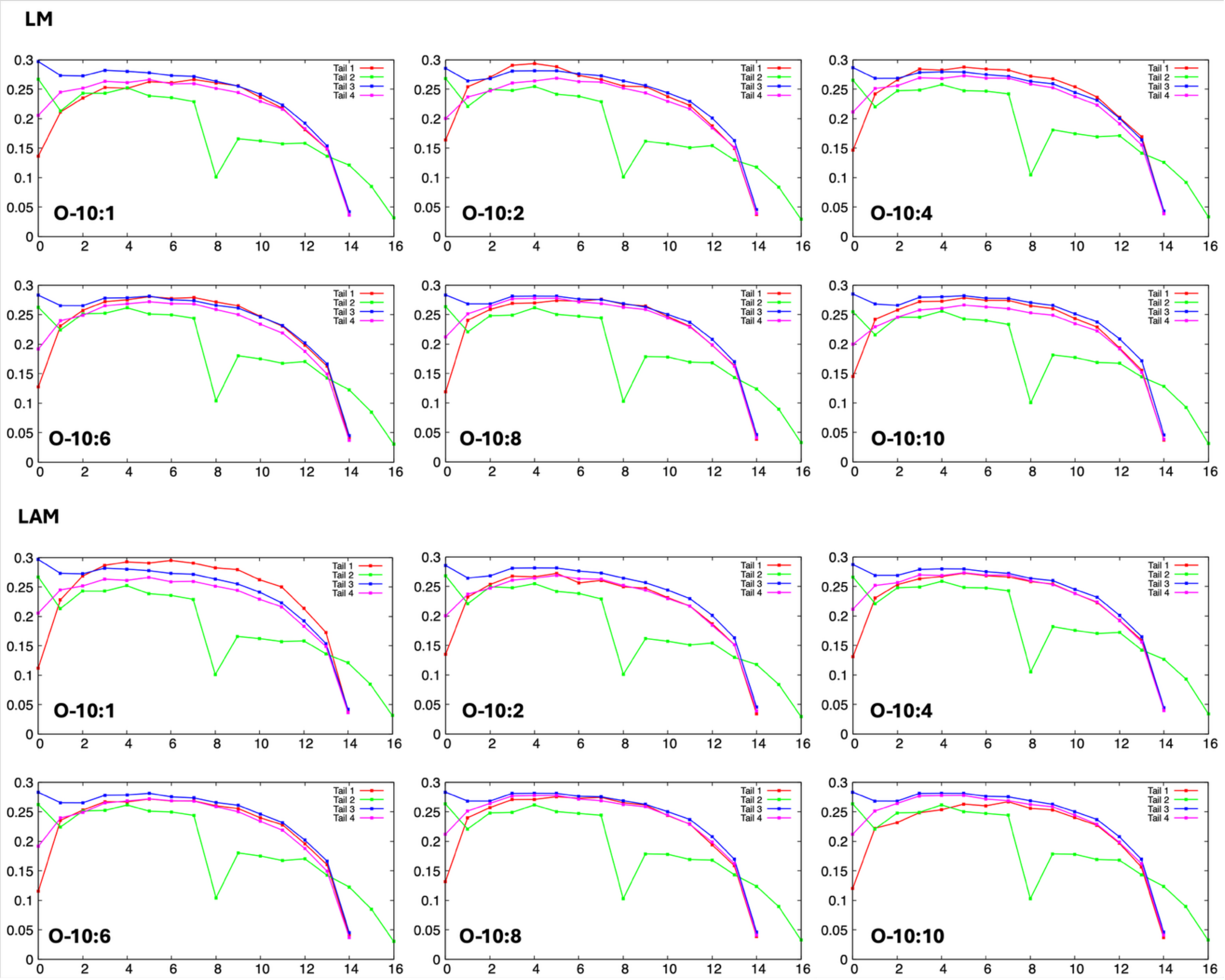


**Figure S4.** Outer leaflet symmetric order parameters for each carbon in LM and LAM acyl chains in **Figure S3**.
